# Genetic characterisation of the *Theileria annulata* cytochrome b locus and its impact on buparvaquone resistance in ruminants

**DOI:** 10.1101/2022.05.25.493409

**Authors:** Qasim Ali, Osama Zahid, Moez Mhadhbi, Ben Jones, Mohamed Aziz Darghouth, George Raynes, Kiran Afshan, Richard Birtles, Neil D. Sargison, Martha Betson, Umer Chaudhry

**Affiliations:** Faculty of Veterinary and Animal Sciences, University of Agriculture, Dera Ghazi Khan, Pakistan; Royal (Dick) School of Veterinary Studies, University of Edinburgh, UK; School of Veterinary Medicine, University of Surrey, UK; Department of Zoology, Faculty of Biological Sciences, Quaid-i-Azam University, Islamabad, Pakistan; Laboratoire de Parasitologie, École Nationale de Médecine Vétérinaire, Université de la Manouba, Sidi Thabet, Tunisia; School of Science, Engineering and Environment, University of Salford, UK

**Keywords:** Buparvaquone, *Theilieria annulata*, resistance, soft selective sweep, hard selective sweep

## Abstract

Control of tropical theileriosis depends on the use of a single drug, buparvaquone, the efficacy of which is compromised by the emergence of resistance. The present study was undertaken to improve understanding of the role of mutations conferring buparvaquone resistance in *Theileria annulata*, and the effects of selection pressures on their emergence and spread. First, we investigated genetic characteristics of the cytochrome b locus associated with buparvaquone resistance in 10 susceptible and 7 resistant *T. annulata* isolates. The 129G (GGC) mutation was found in the Q_01_ binding pocket and 253S (TCT) and 262S (TCA) mutations were identified within the Q_02_ binding pocket. Next, we examined field isolates and identified cytochrome b mutations 129G (GGC), 253S (TCT) and 262S (TCA) in 21/75 buffalo-derived and 19/119 cattle-derived *T. annulata* isolates, providing evidence of positive selection pressure. Both hard and soft selective sweeps were identified, with striking differences between isolates. For example, 19 buffalo-derived and 7 cattle-derived isolates contained 129G (GGC) and 253S (TCT) resistance haplotypes at a high frequency, implying the emergence of resistance by a single mutation. Two buffalo-derived and 11 cattle-derived isolates contained equally high frequencies of 129G (GGC), 253S (TCT), 129G (GGC)/253S (TCT) and 262S (TCA) resistance haplotypes, implying the emergence of resistance by pre-existing and or recurrent mutations. Phylogenetic analysis further revealed that 9 and 21 unique haplotypes in buffalo and cattle-derived isolates were present in a single lineage, suggesting a single origin. We propose that animal migration between farms is an important factor in the spread of buparvaquone resistance in endemic regions of Pakistan. The overall outcomes will be useful in understanding how drug resistance emerges and spreads, and this information will help design strategies to optimise the use and lifespan of the single most drug use to control tropical theileriosis.

## 1 Introduction

Tropical theileriosis, caused by *Theileria annulata* and transmitted by *Hyalomma* ticks, is one of the most important livestock diseases in Asia and North Africa. The disease is highly pathogenic, infecting mononuclear cells of mammalian hosts (Nene et al., 2016). *T. annulata* has a major impact on food production in low- and middle-income countries throughout South Asia, the Middle East and North Africa, where efficient agriculture and livestock production is a priority under the UN Sustainable Development Goals. The disease causes severe economic losses due to high mortality and morbidity of livestock, having a significant impact on meat and milk production (Jabbar et al., 2015).

Hydroxynaphthoquinone drugs, including buparvaquone and atovaquone compounds, are widely used for the treatment of protozoan parasites of livestock and humans, respectively. These drugs bind to parasite cytochrome b, inhibiting mitochondrial respiration (Fry and Pudney, 1992; McHardy et al., 1983; McHardy and Morgan, 1985; McHardy et al., 1985). Different studies have shown associations between resistance to atovaquone in *Plasmodium* and *Toxoplasma* species and mutations in cytochrome b catalytic Q_01_ (codon 129-148), and oxidative Q_02_ (codon 244-266) binding pockets (Kessl et al., 2005; Kessl et al., 2006; Kessl et al., 2007; Korsinczky et al., 2000; Schwobel et al., 2003; Song et al., 2015; Srivastava et al., 1999). Atovaquone resistance-associated cytochrome b mutations at codon 268 have been reported in *Plasmodium falciparum* (David et al., 2003; Farnert et al., 2003; Fivelman et al., 2002; Musset et al., 2006; Schwartz et al., 2003); mutations at codon 133, 144 and 284 in *Plasmodium berghei* (Siregar et al., 2008; Syafruddin et al., 1999); and mutations at codon 129 and 254 in *Toxoplasma gondii* (McFadden et al., 2000).

The control of theileriosis heavily depends on the use of buparvaquone; it is the only effective commercially available drug for use in livestock. Clinical cases of buparvaquone resistance have been reported and represent a serious threat to efficient livestock production (Cui et al., 2015; Mhadhbi et al., 2010; Muraguri et al., 2006). Current understanding of the mechanism of buparvaquone resistance in *T. annulata* is limited, albeit links between mutations in cytochrome b and the resistance phenotype have been demonstrated (Mhadhbi et al., 2015). Moreover, a few studies have demonstrated buparvaquone resistance-associated mutations at codon 129, 253 and 262 in *T. annulata* isolates collected from Iran (Sharifiyazdi et al., 2012), Tunisia (Mhadhbi et al., 2015) and Sudan (Chatanga et al., 2019).

Selective sweep models could potentially explain how resistance mutations emerge in parasites; whereby the use of antiparasitic drugs provides positive selection pressure for adaptive mutations in the resistance candidate loci of the parasite (Chaudhry and Gilleard, 2015). A hard selective sweep is characterised by a single resistance haplotype rising from a recent mutation at a high frequency in each parasite isolate (Chaudhry et al., 2020; Chaudhry et al., 2015; Shaukat et al., 2019). A soft selective sweep is characterised by the presence of multiple resistance haplotypes at a high frequency in each isolate, derived from either recurrent mutations appearing after the onset of selection, or pre-existing mutations before the onset of selection (Chaudhry et al., 2016a; Chaudhry et al., 2020; Shaukat et al., 2019). Understanding adaptive mutations in response to selection can help to determine their origin and spread (Chaudhry and Gilleard, 2015). For example, phylogenetic models might imply that new resistance mutations could arise from a single origin, become fixed by selection and then spread through parasites by migration. This scenario would likely be a consequence of the gene flow of drug resistance alleles through host movement. In contrast, resistance mutations could repeatedly arise from multiple origins, become fixed by selection and migrate between parasites as a result of host movement (Chaudhry et al., 2016a; Chaudhry et al., 2015; Shaukat et al., 2019; Shaukat et al., 2021).

High throughput deep amplicon sequencing using the Illumina Mi-Seq platform is an accurate and relatively low-cost method for the identification of amplicon sequence variants in parasites. The method has transformed the study of benzimidazole resistance in nematode parasites (Ali et al., 2019; Sargison et al., 2019), and pyrimethamine resistance in *Plasmodium* (Shaukat et al., 2019; Shaukat et al., 2021) and has the potential to open new areas of research to improve understanding of buparvaquone resistance in *Theileria*. The method can generate sequence reads up to 600 bp in length by targeting primer binding to conserved sites. Moreover, with the use of barcoded primers, up to 384 samples can be pooled and sequenced in a single Mi-Seq run.

The overall aim of the present study was to develop a genetic approach to improve understanding of the *T. annulata* cytochrome b locus in buparvaquone resistance. This allowed the analysis of alleles conferring resistance and susceptibility to investigate different models of emergence and the spread of associated mutations. The specific objectives were: (i) To characterise single nucleotide polymorphism (SNPs) in the cytochrome b locus and their genetic role in buparvaquone resistance in *T. annulata*. (ii) To investigate the consequences of selection pressure on the emergence and spread of *T. annulata* cytochrome b resistance-conferring mutations in the field isolates. Overall, the results will lead to an increased understanding of buparvaquone resistance in an important tropical livestock pathogen, with the potential to inform enhanced animal health, improved global food security, and poverty alleviation through reduced production losses.

## 2 Materials and methods

### 2.1. Buparvaquone susceptible and resistant T. annulata isolates

Four infected cell lines (TA-ank, TA-but, TA-has, TA-mor) had been produced by the infection of blood mononuclear cells with laboratory-maintained stocks of *T. annulata* held in the Roslin Institute of the University of Edinburgh, originally isolated in Turkey, India, Tunisia, and Morocco (Katzer et al., 1994). The stock was selected to represent buparvaquone susceptible isolates. Genomic DNA of was prepared using 1,000 μl Direct PCR lysis reagent (Viagen), 50 μl proteinase K solution (Qiagen), and 50 μl 1M dithiothreitol (DTT). To extract the gDNA, 50 μl of each isolate was transferred into a fresh tube and centrifuged for 5 minutes, before removing the supernatant and mixing with 25 μl lysis buffer (Viagen), Proteinase K (New England BioLabs) and dithiothreitol (DTT).

Six known buparvaquone susceptible isolates (674, Battan C, Battan P4, Chargui P5, Jed4, Jed4 P10) were selected from different endemic regions in the north-east of Tunisia. Battan C, Battan P4, Jed4 P10 and Jed4 isolates were chosen based on previous studies confirming the susceptibility of these stocks to buparvaquone (Darghouth et al., 1996). The Chergui P5 and 674 isolates were chosen from the animals cured after the first injection of buparvaquone. Seven known buparvaquone resistant isolates (ST2/13, ST2/19, 739, 881III, 5911, 8307, BC2T) were selected from different endemic regions in the north-east of Tunisia. Two buparvaquone resistant isolates (ST2/13, ST2/19) were taken from the previous study by Mhadhbi *et al* (2015). Resistant stocks ST2/13 and ST2/19 were isolated from a clinical case of treatment failure against tropical theileriosis using buparvaquone. The samples were collected at different times after treatment as follows: isolate ST2/13 was taken 24 hours after the first treatment and the ST2/19 isolate was taken 48 hours after the third treatment (Mhadhbi et al., 2015). Five stocks (739, 881III, 5911, 8307, BC2T) were initially isolated at various times points from tropical theileriosis clinical cases of treatment failure based on the absence of clinical improvement despite the early initiation of treatment with the conventional dose of buparvaquone, and the persistence of a high parasitaemia after at least three doses of buparvaquone. All isolates were collected by lymph node punctures or from whole blood and cultured in a complete RPMI 1640 medium. They had been used at the low passage to avoid *in-vitro* pressure selection (Mhadhbi et al., 2015). Genomic DNA was extracted using a Promega Wizard DNA Purification kit (Madison, WI, USA) according to the manufacturer’s instructions.

### 2.2. Theileria annulata field isolates

Field isolates were taken from suspected piroplasm infected cattle and buffalo presented at veterinary clinics across the endemic regions of Pakistan between 2020 and 2021. Moreover, piroplasm-negative cattle blood samples were provided by Dr Tim Connelly, Roslin Institute, University of Edinburgh. The samples were collected by para-veterinary staff under the supervision of local veterinarians following consent from the animal owners. The study was approved by the Institutional Review Board of the Quaid-i-Azam University Islamabad Pakistan (No. #BEC-FBS-QAU2017). Peripheral blood smears were prepared and stained with 4% Giemsa and examined microscopically to detect piroplasms. Genomic DNA was isolated from positive samples by lysis with GS buffer containing proteinase K as described in the TIANamp Blood DNA Kit (TIANGEN Biotech Co. Ltd, Beijing) and stored at -20°C. ‘Haemoprotobiome’ high-throughput sequencing was performed on piroplasm-positive blood samples to confirm the presence of *T. annulata* as previously described (Chaudhry et al., 2019). Briefly, 194-piroplasm positive field samples [buffalo (n=75), cattle (n=119)] were examined to identify the species of *T. annulata* involved in the infections.

### 2.3. PCR amplification, Illumina Mi-Seq run and bioinformatics data handling of cytochrome b locus

A 516 bp fragment of the cytochrome b locus of *T. annulata* laboratory and field isolates was amplified. The primer sets, adapter/barcoded PCR amplification conditions and magnetic bead purification were previously described by Chaudhry et al. (2021) (Supplementary Table S1 and S2). Ten μl of each barcoded bead purified PCR product were combined to make a pooled library which was subject to agarose gel electrophoresis. Cytochrome b products were excised and purified from the gel using a commercial kit (QIAquick Gel Extraction Kit, Qiagen, Germany), followed by purification of eluted DNA using AMPure XP Magnetic Beads (1X) (Beckman Coulter, Inc.). Purified products were then combined into a single purified DNA pool library. The library was quantified with qPCR library quantification kit (KAPA Biosystems, USA) and then run on an Illumina MiSeq Sequencer using a 600-cycle pair-end reagent kit (MiSeq Reagent Kits v2, MS-103-2003) at a concentration of 15nM with the addition of 15% Phix Control v3 (Illumina, FC-11-2003) described by Chaudhry et al. (2021).

The Illumina Mi-Seq post-run processing uses the barcoded indices to separate all sequences by sample and generate FASTQ files. The FASTQ files of the buparvaquone susceptible and resistant *T. annulata* isolates and *T. annulata* field isolates have been made freely available through the Mendeley database (doi: 10.17632/n6xpdcntvf.1). These files were analysed using Mothur v1.39.5 software (Kozich et al., 2013; Schloss et al., 2009) with modifications in the standard operating procedures of Illumina Mi-Seq in the Command Prompt pipeline described by Chaudhry et al. (2021).

### 2.4. Allele frequency of buparvaquone resistance-associated SNPs in the cytochrome b locus

The calculation of the relative allele frequencies of *T. annulata* cytochrome b resistance-associated mutations identified in the buparvaquone susceptible and resistant isolates and field isolates was performed by dividing the number of sequence reads of each isolate by the total number of reads (R Core Team, 2013; package ggplot2).

### 2.5. Genetic models of buparvaquone resistance-associated SNPs in the cytochrome b locus

The sequence reads of the *T. annulata* cytochrome b locus from each isolate were imported into the FaBox 1.5 tool (birc.au.dk) to collapse the sequences that showed 100% base pair similarity after corrections into a single haplotype (freely available through the Mendeley database doi: 10.17632/n6xpdcntvf.1). The haplotypes of cytochrome b were then aligned using the MUSCLE alignment tool in Geneious v10.2.5 software (Biomatters Ltd, New Zealand) for the analysis of the emergence of buparvaquone resistance-associated mutations in the cytochrome b locus. The haplotypes of the *T. annulata* cytochrome b locus were selected for neutrality analysis. Tests for selective neutrality were analysed to determine whether the observed frequency distribution of sequence polymorphism in *T. annulata* cytochrome b locus departed from neutral expectations using the DnaSP 5.10 software package (Librado and Rozas, 2009). The neutrality tests included Tajima’s D (Tajima, 1989) and Fay & Wu’s H (Fay and Wu, 2000) methods.

A split tree of cytochrome b haplotypes was created in the SplitTrees4 software (bio-soft.net) using the neighbour-joining method and the JukesCantor model of substitution. The appropriate model of nucleotide substitutions for neighbour-joining analysis was selected by using the jModeltest 12.2.0 program (Posada, 2008). The network tree of *T. annulata* cytochrome b haplotypes was produced based on the neighbour-joining algorithm built on a sparse network and the epsilon parameter is set to zero default in Network 4.6.1 software (Fluxus Technology Ltd.). All unnecessary median vectors and links were removed with the star contractions. The number of mutations separating adjacent sequence nodes was indicated along connecting branches and the length of the lines connecting the haplotypes is proportional to the number of nucleotide changes.

## 3. Results

### 3.1. Buparvaquone resistance-associated SNPs in susceptible and resistant T. annulata isolates

A 516 bp region of the cytochrome b locus corresponding to the buparvaquone binding pockets [(Q_01_ (codon 129-148) and Q_02_ (codon 244-266)] of the susceptible and resistant *T. annulata* isolates was selected for deep amplicon sequencing. The sequences of 10 susceptible and 7 resistant isolates were compared. Eleven point mutations in the *T. annulata* cytochrome b locus were found at codons A133V (GCT/GTT), V135F (GTC/TTC), I136L (ATA/CTA), G138V (GGT/GTT), L139F (TTG/TTT), L140F (TTG/TTT), K141N (AAA/AAC), F143L (GGA/GTA), G144V (GGA/GTA) and T146A (ACT/GCT), within the cytochrome b Q_01_ binding pocket. These SNPs were present at similar frequencies across the buparvaquone susceptible and resistant *T. annulata* isolates (Table 1). One point mutation was found in the Q_01_ binding pocket at codon 129G (GGC) in resistant isolates compared to S129 (AGC) in susceptible isolates (Table 1). Similarly, two-point mutations were found within the Q_02_ binding pocket at codons 253S (TCT) and 262S (TCA) in resistant isolates compared to P253 (CCT) and L262 (TTA) in susceptible isolates (Table 1). Overall, the results provide a comprehensive analysis of cytochrome b mutations potentially involved in the development of buparvaquone resistance and the first genetic link between mutations and drug resistance in *T. annulata*.

**Table 1.**
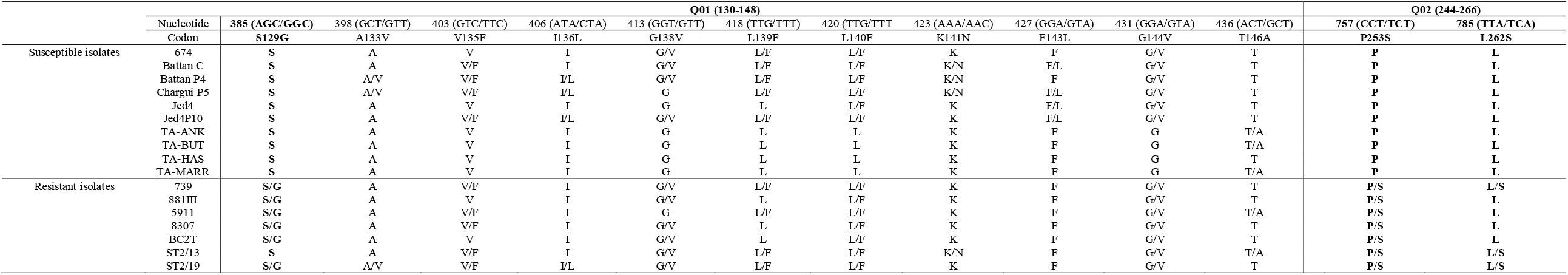
Cytochrome b mutations identified in buparvaquone susceptible *T. annulata* isolates collected from an endemic region of Turkey, India, Tunisia, and Morocco and resistant *T. annulata* isolates collected from an endemic region of Tunisia. Comparison shows the buparvaquone susceptible (674, Battan C, Battan P4, Chargui P5, Jed4, Jes4P10, TA-ANK, TA-BUT, TA-HAS, TA-MARR) and resistant isolates (739, 881III, 5911, 8307, BC2T, ST2/13, ST2/19). Nonsynonymous mutations at positions S129G (AGC/GGC), P253S (CCT/TCT) and L262S (TTA/TCA) in the binding pockets Q_01_ and Q_02_ of the cytochrome b locus differ between susceptible and resistant *T. annulata* isolates.

### 3.2. Buparvaquone resistance-associated SNPs in buffalo- and cattle-derived T. annulata field isolates and their impact on positive selection pressure

A 516 bp fragment of the *T. annulata* cytochrome b locus was amplified from 75 buffalo-derived and 119 cattle-derived isolates across the endemic regions of Pakistan (Supplementary Table S3). The relative allele frequencies of three mutations S129G (AGC/GGC), P253S (CCT/TCT) and L262S (TTA/TCA) correspond to the buparvaquone binding pockets were determined. Buparvaquone resistance-associated mutations 129G (GGC), 253S (TCT) and 129G (GGC)/253S (TCT) were present in 21/75 buffalo-derived *T. annulata* (Fig. 1, Table 2). The 129G (GGC) mutation was present at frequencies of 4.3-100%. The 253S (TCT) mutation was present at frequencies of 19.2-27.3%. Conversely, 129G (GGC)/253S (TCT) double mutations were detected at frequencies of 1.3-4.3%. 262S (TCA) was not detected in any buffalo-derived *T. annulata* isolates (Fig. 1, Table 2). In contrast, buparvaquone resistance-associated mutations 129G (GGC), 253S (TCT) and 262S (TCA) were present in 19/119 cattle-derived *T. annulata* isolates (Fig. 1, Table 2). The 129G (GGC) mutation was present at frequencies of 1.4-100%. 253S (TCT) mutation was present at frequencies between 3.9-43%. 262S (TCA) mutation was detected at frequencies of 2.6-13.7%. 129G (GGC)/253S (TCT) was not detected in any cattle-derived *T. annulata* (Fig. 1, Table 2). Overall, the results indicated the positive selection pressure on the cytochrome b locus of 21 buffalo-derived and 19 cattle-derived *T. annulata* isolates.

**Table 2.**
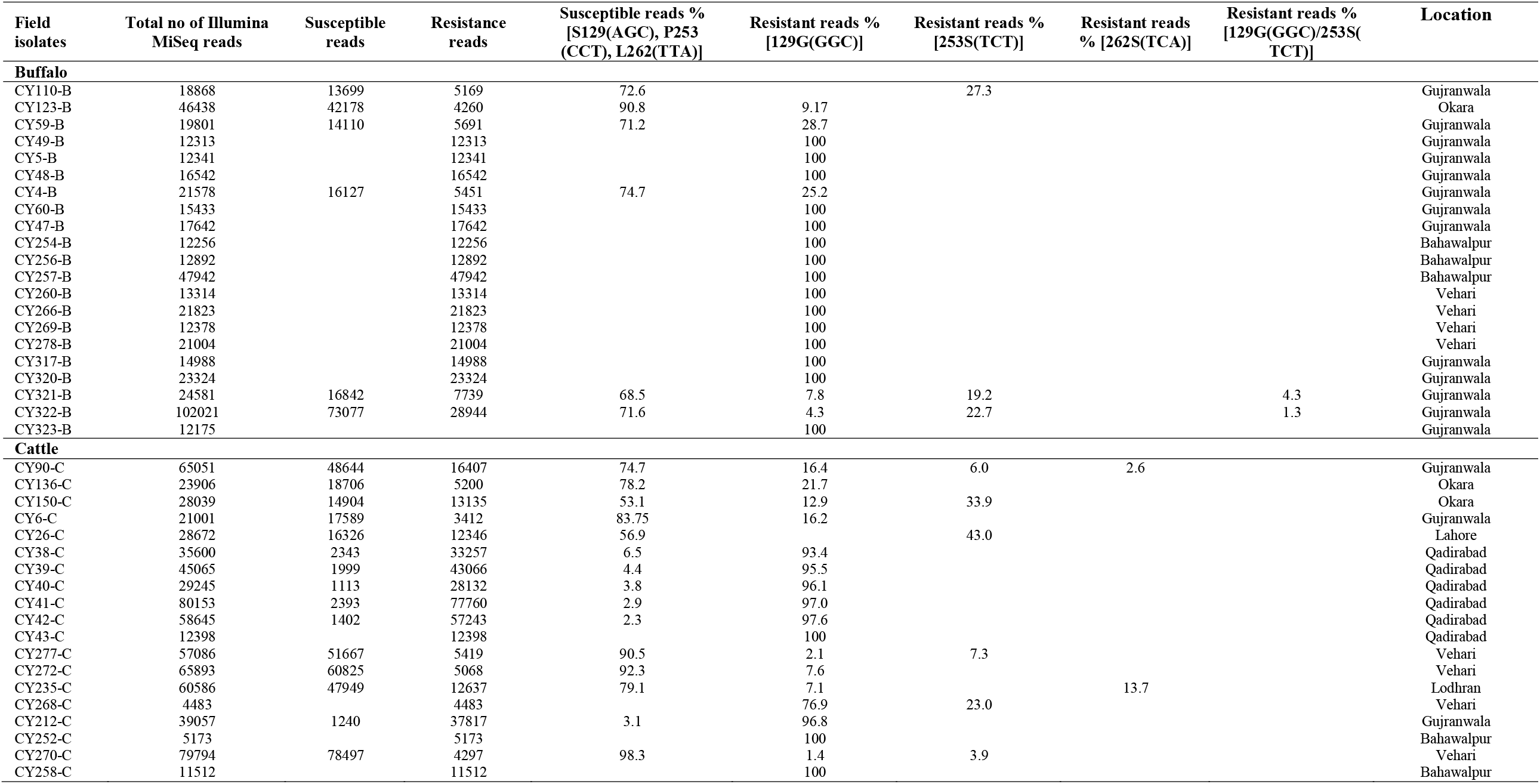
Relative allele frequencies of the cytochrome b mutations in 21 buffalo-derived and 19 cattle-derived *T. annulata* resistant isolates collected from the endemic region of Pakistan. Each isolate was characterized by susceptible [S129 (AGC), P253 (CCT) and L262 (TTA)] and resistance mutations [129G (GGC), 253S (TCT), 262S (TCA) and 129G (GGC)/253S (TCT)]. The relative allele frequency of the resistant versus susceptible mutations was based on the SNPs identified using the deep amplicon sequencing method.

**Figure 1:**
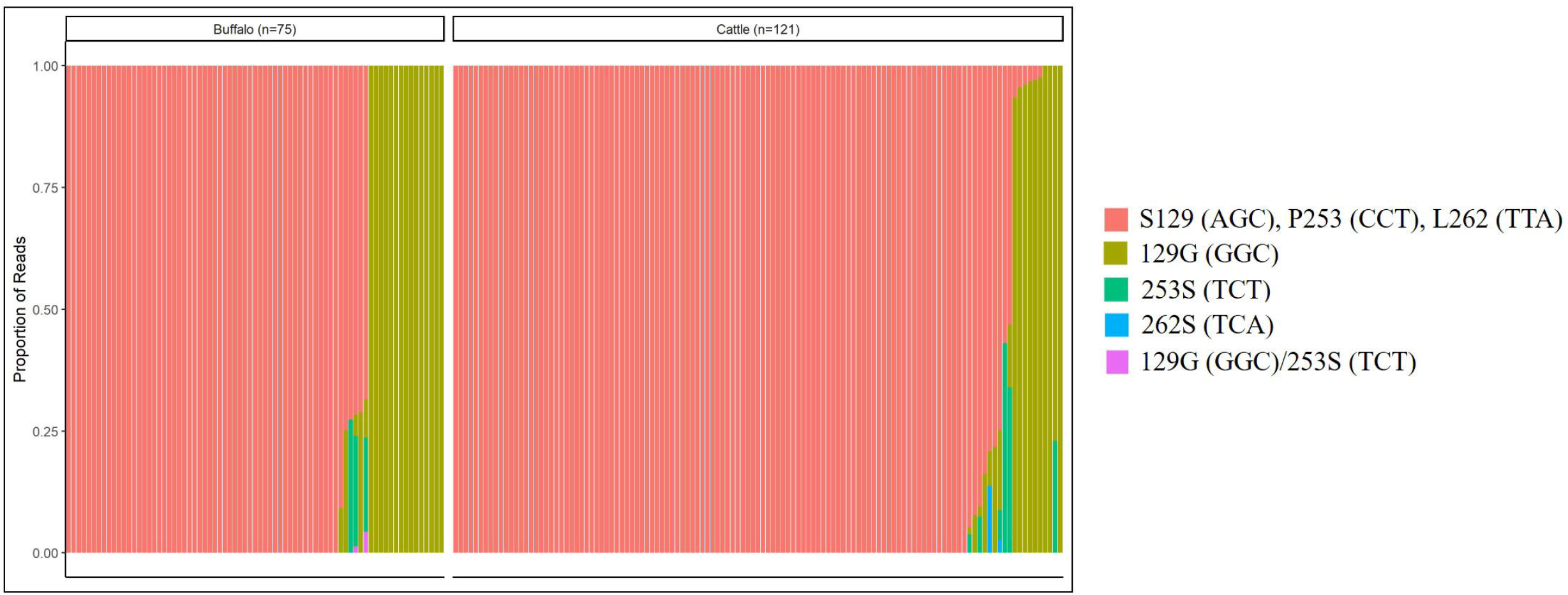
Relative allele frequencies of cytochrome b resistance-associated mutations in 75 buffalo-derived and 119 cattle-derived *T. annulata* isolates collected from endemic regions of Pakistan. Each isolate shows the relative frequency of the resistant versus susceptible SNP based on allele quantification by deep amplicon sequencing (Supplementary Table S3). The susceptible SNP frequency - S129 (AG), P253 (CCT) and L262 (TTA) - is shown in red and resistance-associated SNP frequency - 129G (GGC), 253S (TCT), 262S (TCA) and 129G (GGC)/ 253S (TCT) - is shown in light green, dark green, blue and pink.

### 3.3. Nature of selective sweeps associated with the emergence of cytochrome b resistance-associated SNPs in Buffalo- and cattle-derived T. annulata field isolates

The analysis of 21 buffalo-derived *T. annulata* revealed that 18 isolates contained the 129G (GGC) resistance haplotype and one isolate (CY110-B) contained the 253S (TCT) resistance haplotype at a high frequency of 100% (Fig. 2A, Table 3), providing evidence of hard selective sweep patterns. In contrast, the CY321-B isolate contained resistance haplotypes of 129G (GGC) at a frequency of 24.5%, 253S (TCT) at a frequency of 59.9% and 129G (GGC)/253S (TCT) at a frequency of 13.8% (Fig. 2A, Table 3). The CY322-B isolate contained resistance haplotypes of 129G (GGC) at a frequency of 14.4%, 253S (TCT) at a frequency of 79.3% and 129G (GGC)/253S (TCT) at a frequency of 4.6% (Fig. 2A, Table 3). These two isolates had equally high frequencies of cytochrome b resistance-conferring haplotypes demonstrating evidence of soft selective sweep patterns. Neutrality analysis further revealed a significant departure of neutrality in those 2 isolates providing evidence of a signature of selection at the *T. annulata* cytochrome b locus (Table 3).

**Table 3:**
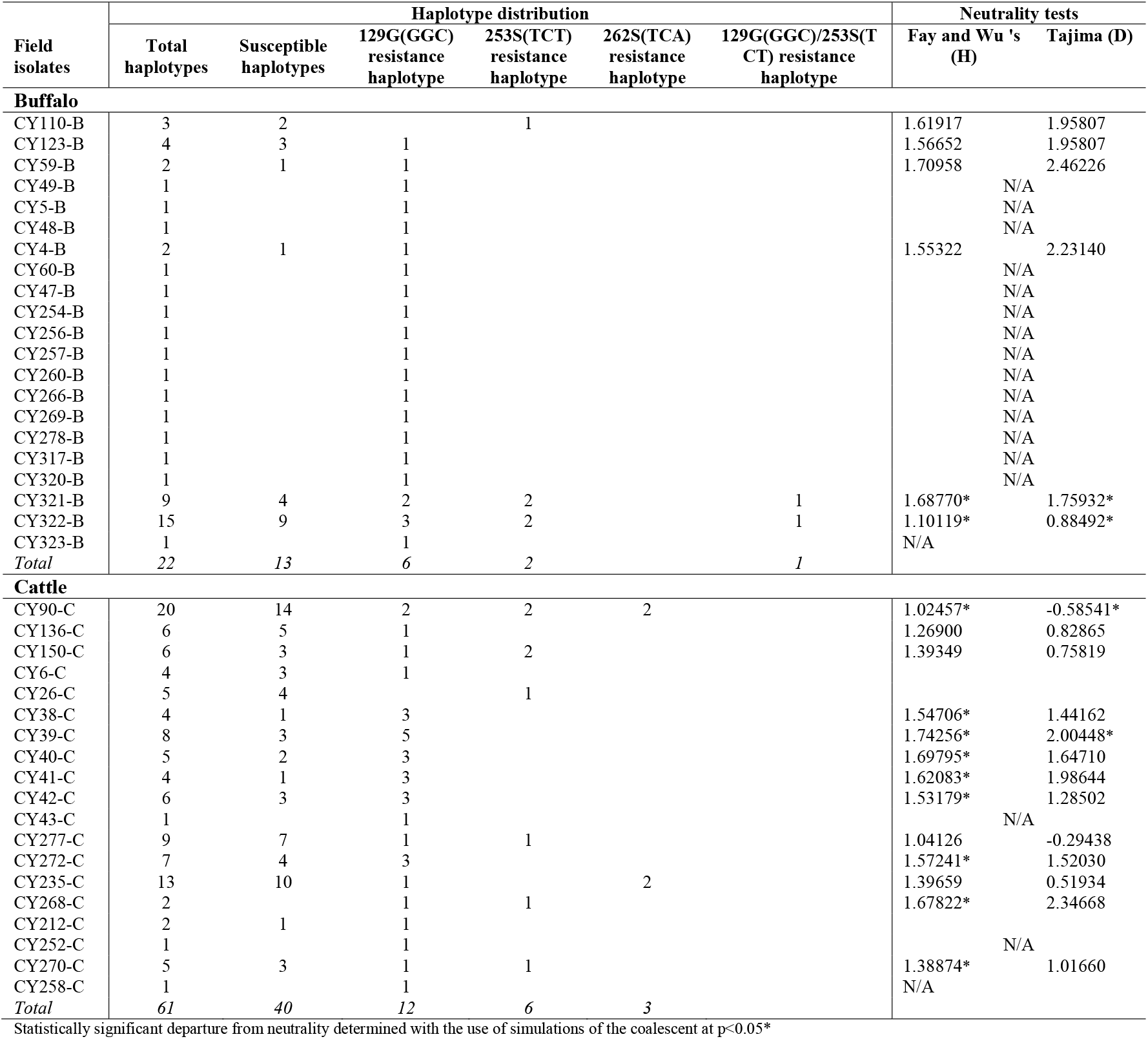
Haplotype diversity and signature of selection at the *T. annulata* cytochrome b locus. A total r of susceptible and resistant haplotypes from 21 buffalo-derived and 19 cattle-derived *T. annulata* isolates the haplotype diversity and signature of selection at the cytochrome b locus.

**Figure 2:**
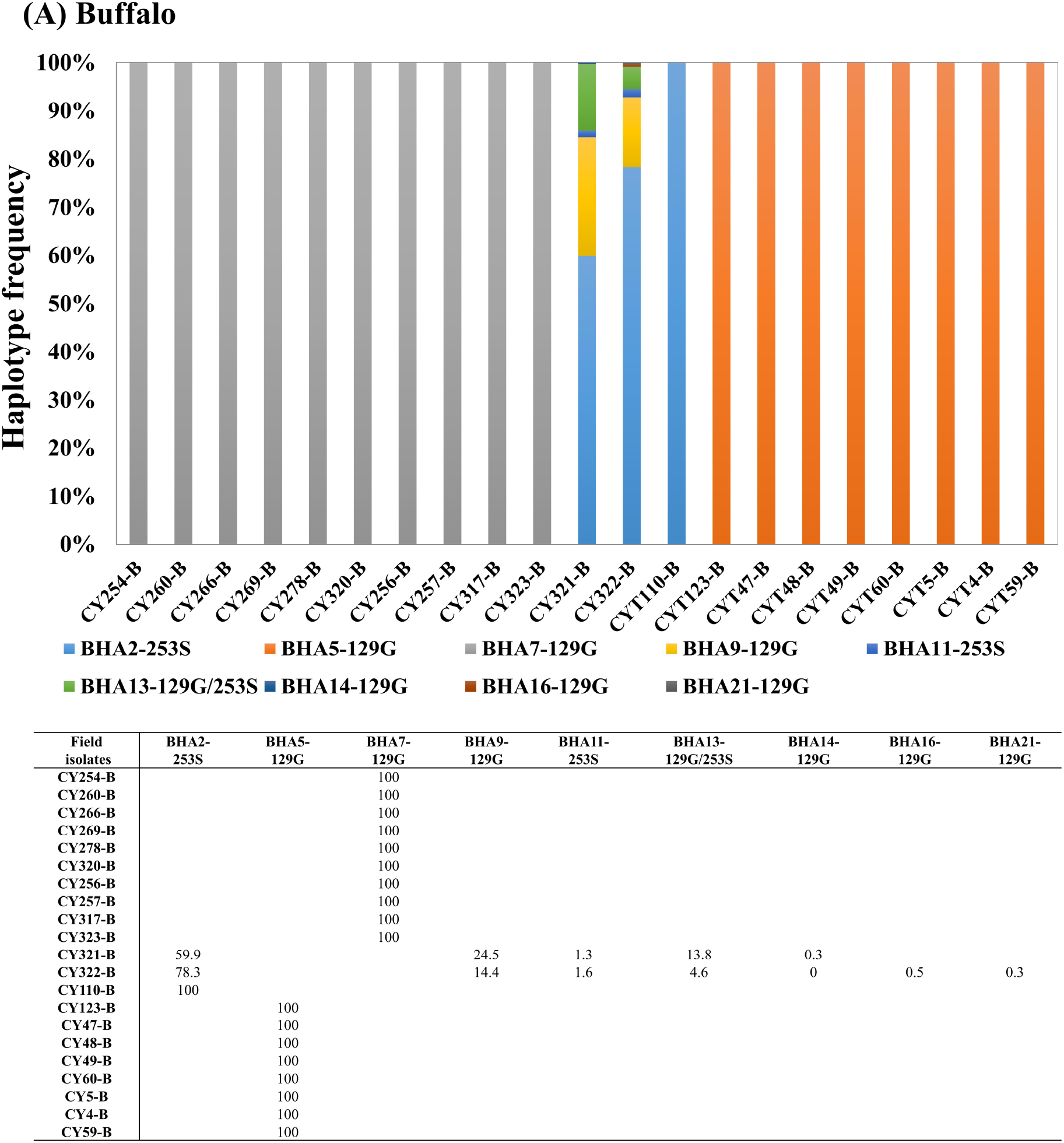
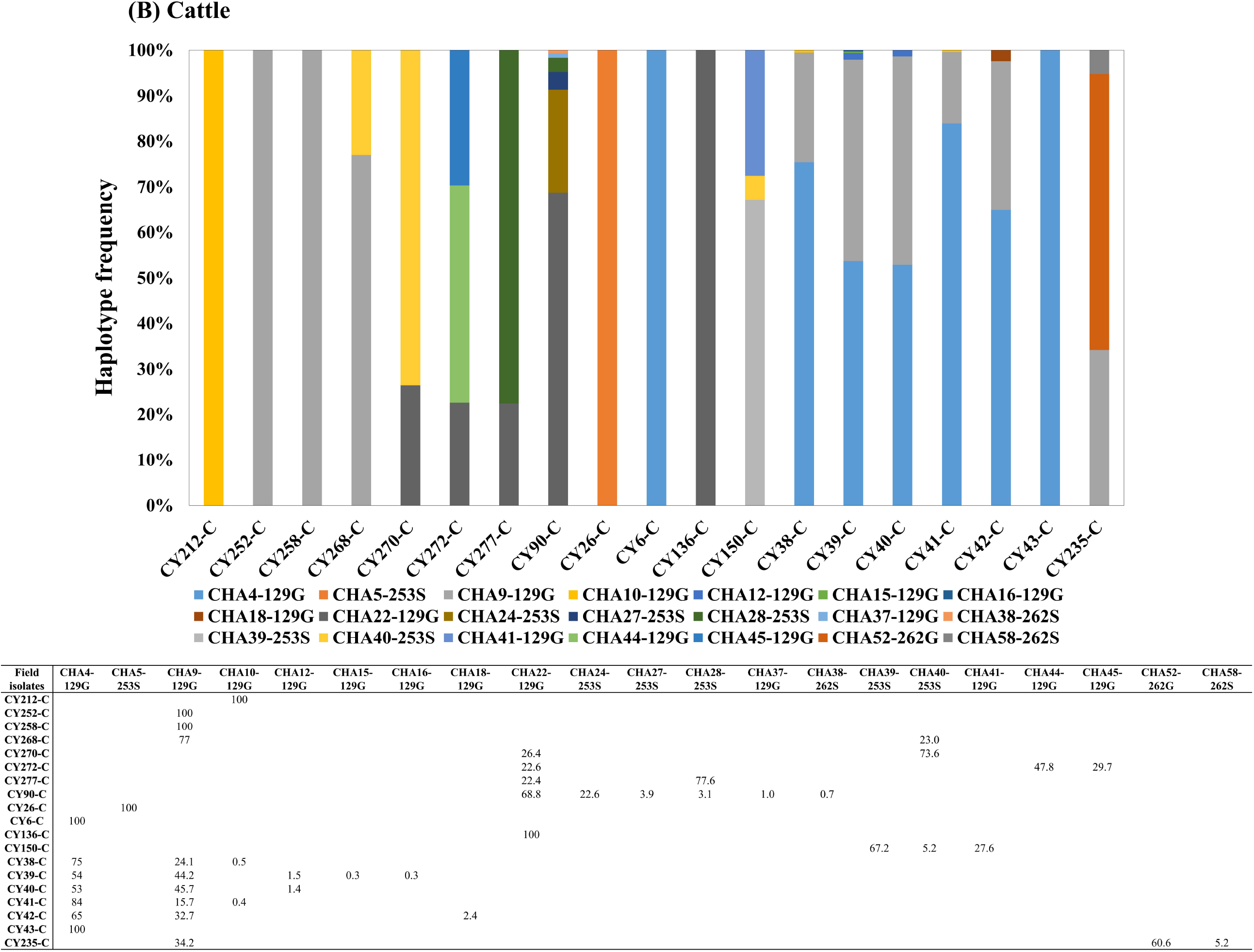
Twenty-two buffalo-derived (2A) and 61 cattle-derived (2B) haplotypes of the cytochrome b locus were generated using the FaBox 1.5 tool (birc.au.dk). The 9/22 buffalo-derived and 21/61 cattle-derived resistance haplotypes (BHA and CHA) are represented by a different colour for the individual isolates (CY). Susceptible haplotypes are not indicated in the figure but are shown in Table 3. The haplotype distribution and the frequency based on the sequence reads generated per *T. annulata* isolates are shown in the insert table.

The analysis of 19 cattle-derived *T. annulata* isolates revealed that 6 (CY212-C, 252-C, 258-C, 6-C, 43-C, 136-C) contained 129G (GGC) resistance haplotypes and one isolate (CY26-C) contained 253S (TCT) resistance haplotype at a high-frequency of 100% (Fig. 2B, Table 3), demonstrating evidence of hard selective sweep patterns. In contrast, 9 isolates (CY268-C, 270-C, 272-C, 277-C, 38-C, 39-C, 40-C, 41-C, 42-C) contained 129G (GGC) resistance haplotypes and one isolate (CY150-C) contained 253S (TCT) resistance haplotypes at frequencies ranging between 16 to 84% (Fig. 2B, Table 3). The CY235-C isolate contained resistance haplotypes of 129G (GGC) at a frequency of 34.2% and 262S (TCA) at a frequency of 60.6%. The CY90-C isolate contained resistance haplotypes of 129G (GGC) at a frequency of 69.8%, 253S (TCT) at a frequency of 22.6% and 262S (TCA) at a frequency of 0.07% (Fig. 2B, Table 3). These 12 isolates had equally high frequencies of cytochrome b resistance-conferring haplotypes demonstrating evidence of soft selective sweep patterns. Neutrality analysis further revealed a significant departure of neutrality in 9 out of 12 isolates providing evidence of a signature of selection at the *T. annulata* cytochrome b locus (Table 3).

### 3.4. Phylogeny of cytochrome b resistance-associated SNPs in buffalo- and cattle-derived T. annulata field isolates and their impact on the origin and spread of buparvaquone resistance

The analysis of 21 buffalo-derived *T. annulata* revealed 22 unique cytochrome b haplotypes. Six haplotypes encoded 129G (GGC) resistance mutations, 2 haplotypes encoded 253S (TCT) resistance mutations, one haplotype encoded 129G (GGC)/253S (TCT) double resistance mutations and 13 haplotypes encoded S129 (AGC), P253 (CCT) and L262 (TTA) susceptible mutations (Fig. 3A, Fig. 4A, Table 3). A split tree analysis showed that 129G (GGC) resistance mutants and 129G (GGC)/253S (TCT) resistance mutants carried 7 haplotypes and 253S (TCT) resistance mutant carried 2 haplotypes were located in a single lineage demonstrating evidence of a single origin. (Fig. 3A). The network analysis showed that the BHA5-129G haplotype was present at a high frequency in eight different isolates collected from the livestock farms in the Okara and Gujranwala regions. The BHA7-129G haplotype was present in ten different isolates collected from the livestock farms in the Okara, Gujranwala, Vehari and Bahawalpur regions. BHA9-129G and BHA13-129G/253S haplotypes were present in two isolates collected from the livestock farms in the Gujranwala region. BHA2-253S haplotype was present in three isolates and BHA11-253S haplotype was present in two isolates collected from the livestock farms in the Gujranwala region (Fig 4A). The data provide evidence of a high level of gene flow, predicted to occur due to the unregulated animal movement. However, three haplotypes (BHA14-129G, BHA16-129G and BHA21-129G) were present at a frequency of 100% in a single isolate collected from the livestock farms in the Gujranwala region (Fig 4A), indicating that the resistance of those haplotypes had not yet spread.

**Figure 3:**
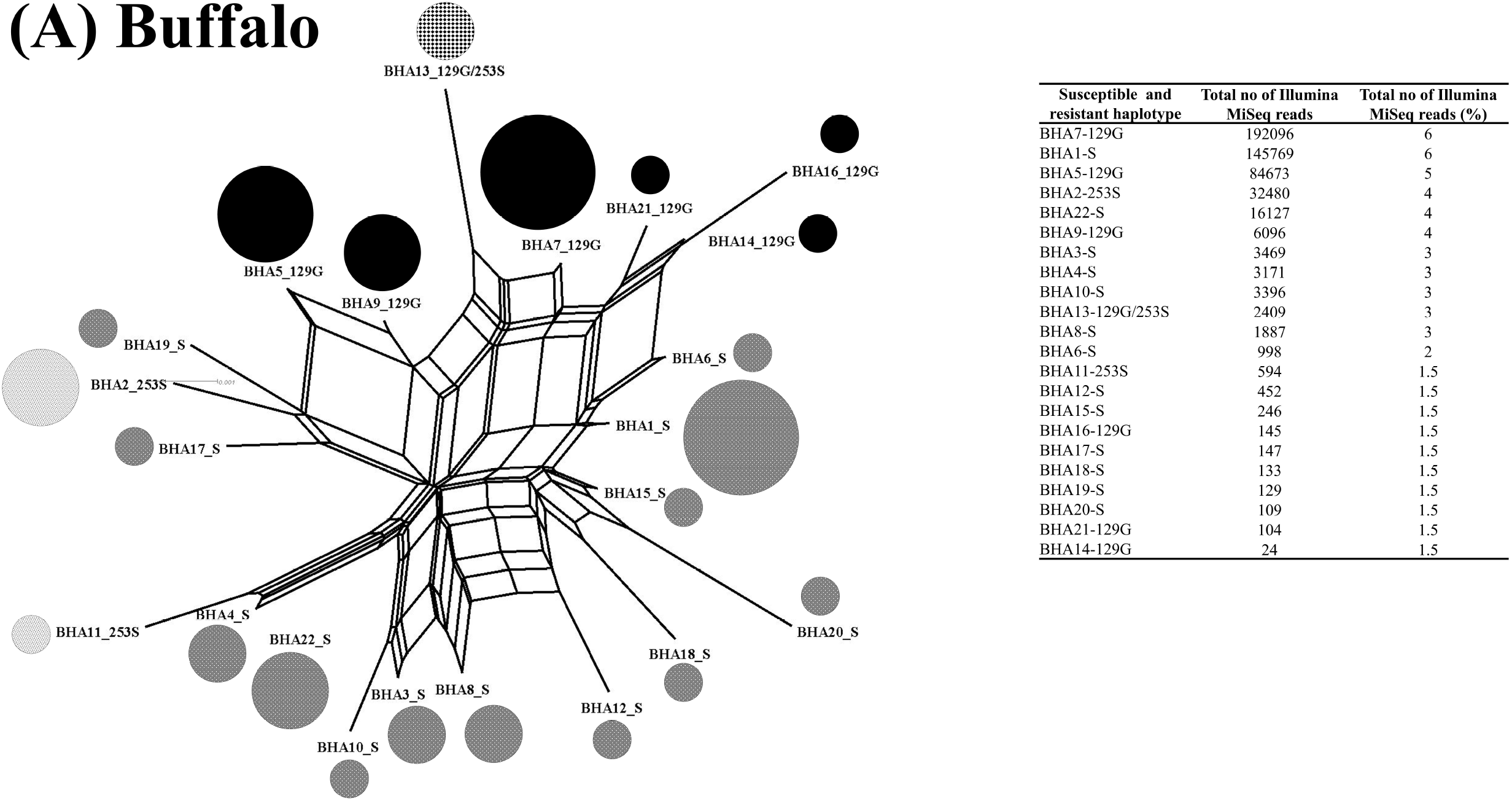
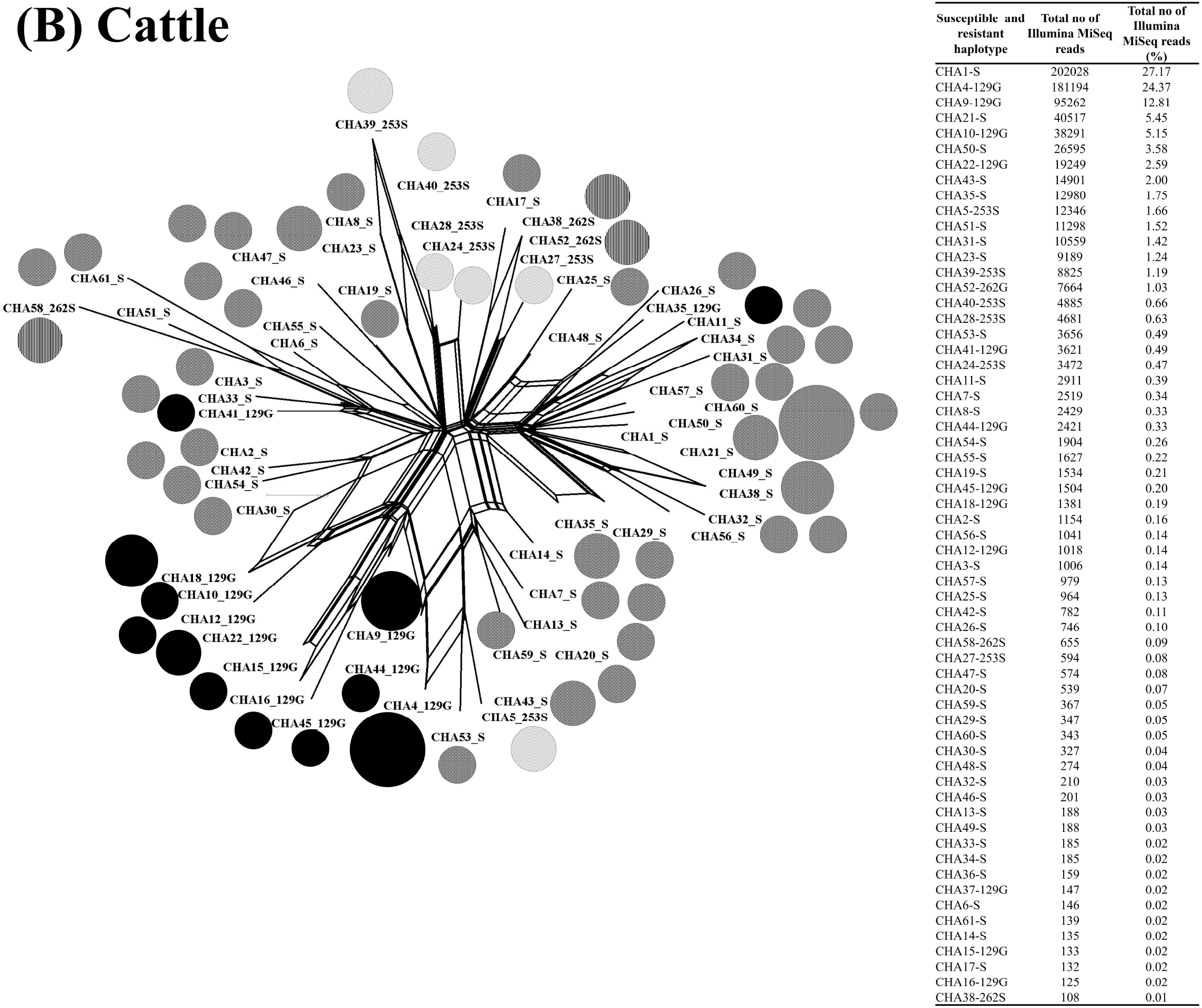
A split tree of 22 buffalo and 61 cattle-derived haplotypes of cytochrome b locus was generated in SplitsTrees4 software (Huson and Bryant, 2006). The pie chart circles represent the different haplotypes and the size of each circle is proportional to the number of sequence reads and frequency generated in that haplotype as indicated in the inserted table. **(A)** The mutation carried buffalo-derived *T. annulata* haplotypes is shaded as follows: thirteen susceptible haplotypes are shown by grey shading; six 129G (GGC) resistant haplotypes are shaded black; one 129G (GGC)/ 253S (TCT) double mutant resistant haplotype is shown by black line shading; and two 253S (TCT) resistant haplotypes are shown by white shading. **(B)** The mutation-carrying cattle-derived *T. annulata* haplotypes are shaded as follows: forty susceptible haplotypes are shown by grey shading; twelve 129G (GGC) resistant haplotypes are shown by black shading; three 262S (TCA) resistant haplotypes are shown by black line shading, and six 253S (TCT) resistant haplotypes are shown by white shading.

**Figure 4:**
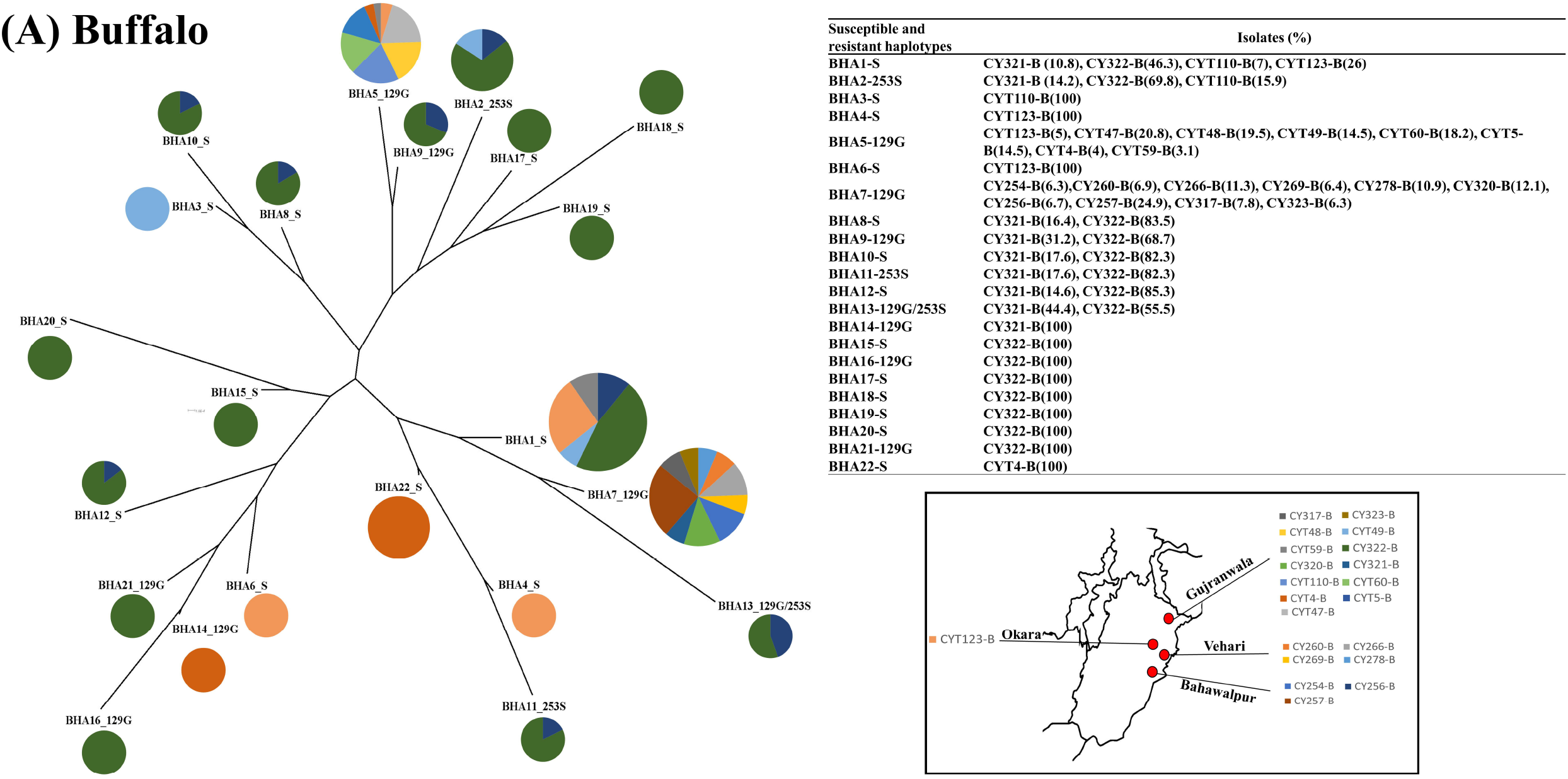
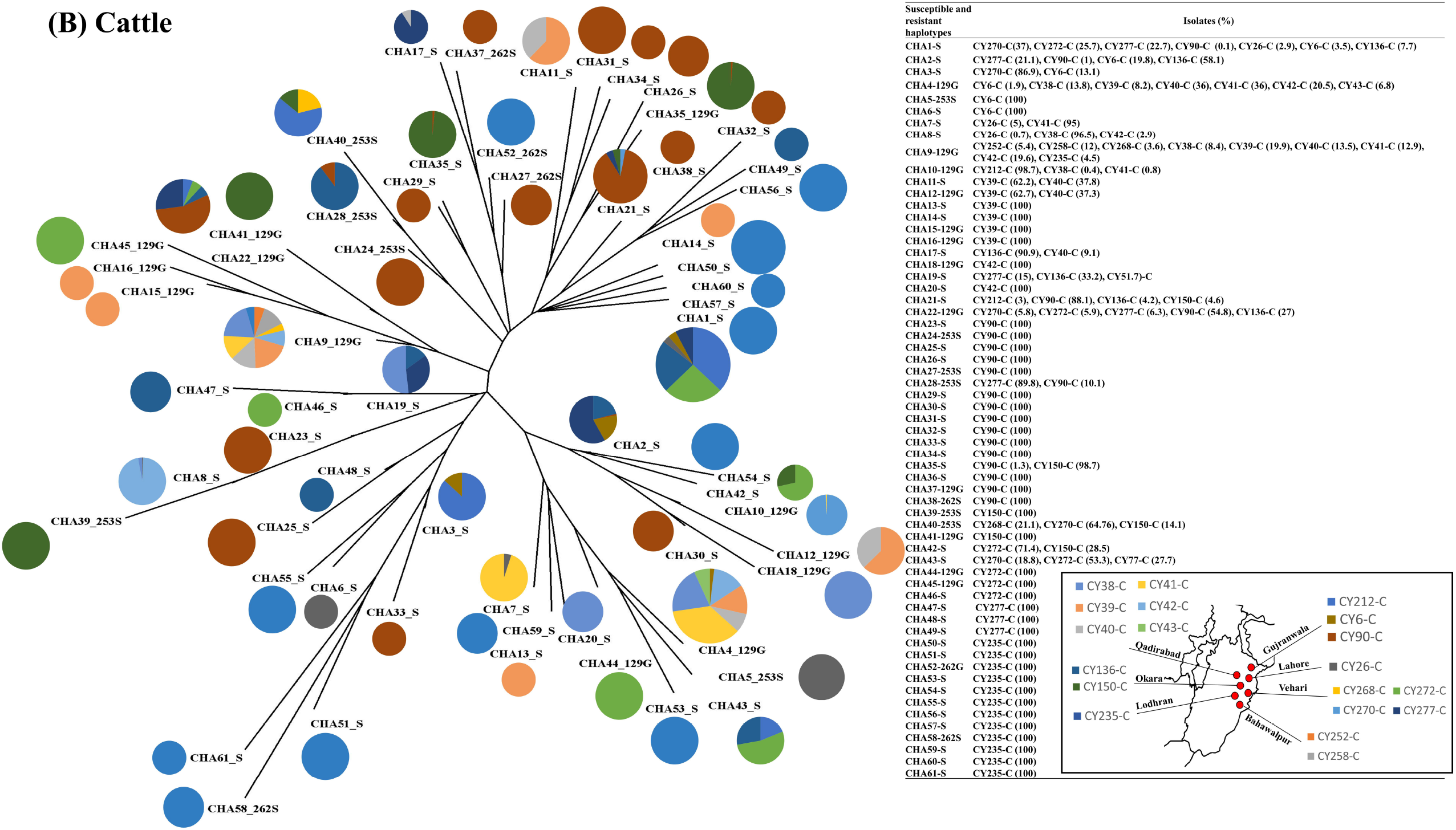
A network tree of 22 buffalo-derived and 61 cattle-derived haplotypes of cytochrome b locus was generated in Network 4.6.1 software (Fluxus Technology Ltd). The size of each pie chart circle representing the haplotype was proportional to the number of sequences generated from different isolates. The colours in the pie chart circles replicate the haplotype frequency and their distribution as indicated in the inserted table. **(A)** The buffalo-derived *T. annulata* carried thirteen susceptible haplotypes, six 129G (GGC) resistant haplotypes, one 129G (GGC)/ 253S (TCT) double mutant resistant haplotype and two 253S (TCT) resistant haplotypes. The mutation-carrying buffalo derived *T. annulata* haplotypes in different livestock farms are shown in the inserted map. **(B)** The cattle-derived *T. annulata* carried forty susceptible haplotypes, twelve 129G (GGC) resistant haplotypes, three 262S (TCA) resistant haplotypes and six 253S (TCT) resistant haplotypes. The mutation-carrying cattle-derived *T. annulata* haplotypes in different livestock farms are shown in the inserted map.

The analysis of 19 cattle-derived *T. annulata* revealed 61 unique cytochrome b haplotypes. 12 haplotypes encoded 129G (GGC) resistance mutations, 6 haplotypes encoded 253S (TCT) resistance mutations, 3 haplotypes encoded 262S (TCA) resistance mutations and 40 haplotypes encoded S129 (AGC), P253 (CCT) and L262 (TTA) susceptible mutations (Fig. 3B, Fig. 4B, Table 3). A split tree analysis showed that 129G (GGC), 253S (TCT) and 262S (TCA) resistance mutants carried 10, 5 and 2 haplotypes were located in a single lineage (Fig. 3B) demonstrating evidence of a single origin (Fig. 3B). The network tree analysis showed that CHA4-129G haplotype was present at a high frequency in 7 different isolates collected from the livestock farms in the Gujranwala and Qadirabad regions. The CHA9-129G haplotype was present in 9 different isolates collected from the livestock farms in the Bahawalpur, Qadirabad, Vehari and Lodhran regions. CHA10-129G haplotype was present in 3 isolates, CHA12-129G in 2 isolates, CHA22-129G in 5 isolates at high frequency collected from the livestock farms in the Gujranwala, Qadirabad, Vehari and Okara regions. The analysis further revealed that CHA28-253S haplotype was present in two different isolates collected from the livestock farms in the Gujranwala and Vehari regions. CHA39-253S haplotype was present in three different isolates collected from the livestock farms in the Vehari and Okara regions (Fig 4B). The data of those 7 haplotypes demonstrate the evidence of a high level of gene flow, predicted to occur due to unregulated animal movement. Seven haplotypes (CHA15-129G, CHA16-129G, CHA18-129G, CHA37-129G, CHA41-129G, CHA44-129G, CHA45-129G) were present at a frequency of 100% in five isolates collected from the livestock farms in the Gujranwala, Qadirabad, Vehari and Okara regions (Fig 4B). Four haplotypes (CHA5-253S, CHA24-253S, CHA27-253S, CHA39-253S) were present at a frequency of 100% in single isolates collected from the livestock farms in the Gujranwala and Okara regions (Fig 4B). CHA38-262S, CHA52-262G, CHA58-262S haplotypes were present at a frequency of 100% in single isolates collected from the livestock farms in the Okara and Lodhran regions (Fig 4B). These data demonstrate that the resistance of these haplotypes was not spread yet.

## 4. Discussion

Buparvaquone has been widely used for the therapy of tropical theileriosis for nearly 30 years. The drug was introduced in the early 1990s and exerts its antiprotozoal actions against *T. annulata*. Buparvaquone resistance in *T. annulata* emerged in 2010 (Mhadhbi et al., 2010). The determinants of resistance to buparavaquone are probably complex in *T. annulata*, evidence suggest that mitochondrial and non-mitochondrial pathways contribute in *T. annulata* resistance to buparvaquone. Indeed, Marsolier et al. 2015 demonstrated that buparvaquone is targeting *Theileria* peptidyl prolyl isomerase to block the signalling pathways leading to the cells proliferations. Interestingly, this enzyme is mutated in a drug resistant isolates (Marsolier et al., 2015). In contrast, the present study is focusing on the mitochondrial pathway for buparvaquone resistance in *T. annulata*. The drug target the cytochrome bc_1_ complex, inhibiting mitochondrial respiration (Fry and Pudney, 1992). The current understanding of the mechanism of resistance is limited, few studies have demonstrated buparvaquone resistance-associated mutations In *T. annulata* isolates (Chatanga et al., 2019; Mhadhbi et al., 2015; Sharifiyazdi et al., 2012). There is a clear need to understand how the resistance mutations against buparvaquone drugs emerge. The frequency with which buparvaquone resistance in *T. annulata* arises and the extent to which it spreads are important considerations for its prevention and control. Therefore, the present study investigates the genetic characteristics of the cytochrome b locus and their impact on the buparvaquone resistance in ruminants. The study further describes genetic models to understand the emergence and spread of buparvaquone resistance at the cytochrome b locus of *T. annulata*.

Deep amplicon sequencing identified fourteen mutations in the cytochrome b catalytic Q_01_ and oxidative Q_02_ binding pockets of *T. annulata* collected from clinical cases of treatment failure in the endemic region of Tunisia. Eleven mutations were present at similar frequencies across the buparvaquone susceptible and resistant *T. annulata* isolates. One mutation at codon 129G (GGC) in resistant isolates and S129 (AGC) in susceptible isolates was found in the Q_01_ binding pocket. Similarly, two mutations at codons 253S (TCT) and 262S (TCA) in resistant isolates and P253 (CCT) and L262 (TTA) in susceptible isolates were found within the Q_02_ binding pocket. Similar studies have been shown that the same mutations in the cytochrome b locus of *T. annulata* isolate collected from clinical cases of treatment failure in Iran (Sharifiyazdi et al., 2012) and Tunisia (Mhadhbi et al., 2015) and Sudan (Chatanga et al., 2019). Overall, the data provide a comprehensive high throughput sequencing analysis of cytochrome b loci involved in the development of buparvaquone resistance and the first genetic link between mutations and drug resistance in *T. annulata*.

Deep amplicon sequencing was then performed to analyse the cytochrome b locus of buffalo- and cattle-derived *T. annulata* field isolates collected from the endemic regions of Pakistan. Buparvaquone resistance-associated mutations 129G (GGC), 253S (TCT) and 129G (GGC)/253S (TCT) were identified in 21 buffalo-derived and 19 cattle-derived *T. annulata* isolates. Overall, buparvaquone resistance-associated SNPs were reported for the first time in buffalo- and cattle-derived *T. annulata* field isolates and their impact on positive selection pressure. Our results are consistent with the study of the pyrimethamine resistance mutations in the protozoan parasite *Plasmodium vivax* (Auliff et al., 2006; Brega et al., 2004; de Pecoulas et al., 1998; Hastings et al., 2005; Imwong et al., 2003; Kaur et al., 2006; Kuesap et al., 2011; Lu et al., 2012; Mint Lekweiry et al., 2012; Ranjitkar et al., 2011; Schunk et al., 2006; Shaukat et al., 2019) and diminazene resistance in the *Trypanosoma brucei* (Carter et al., 1999; Mäser et al., 1999; Matovu et al., 2001). A possible explanation for the differences in the frequency of buparvaquone-conferring mutations with positive selection pressure may be variable drug doses, for example, if the 262S (TCA), 253S (TCT) and 129G (GGC)/253S (TCT) resistance mutations may be selected at low doses of buparvaquone, while 129G (GGC) may occur at higher doses. Overall, the data provide novel information for screening field samples for the detection of buparvaquone resistance mutations for example, in multiple isolates across different geographical regions. The information will aid in understanding how resistance mutations differ between geographical locations with implications for decision-making tools for farmers, farm advisers and veterinarians to design parasite control regimes for individual flocks.

Both hard and soft selective sweep patterns have been demonstrated in the buparvaquone resistance mutations in *T. annulata* in the present study. A single resistance haplotype at a high frequency was detected in the 19 buffalo-derived and 18 cattle-derived isolates. The selective sweeps on these individual isolates were effectively harder with no evidence of a genetic footprint of selection detected by significant departures from the neutrality test. In contrast, multiple resistance haplotypes were detected at high frequencies in the 2 buffalo-derived and 11 cattle-derived isolates. The selective sweep on these individual isolates was effectively softer and a genetic footprint of selection was also detected by significant departures from the neutrality test. The results are consistent with the hypothesis that single and multiple mutations in dhfr locus emerged in the human protozoan parasite *P. vivax* (Shaukat et al., 2019). Overall, the data provide novel information on single and or multiple emergences of resistance mutations in this group of parasites that may have implications for targeted selective treatment, or use of different drug combinations, including new drugs or the modification of current compounds in resistance mitigation strategies.

It will be challenging to determine the emergence of resistance mutations with multiple haplotypes, either due to pre-existing mutations before the onset of positive selection or recurrent mutations after the onset of positive selection (Chaudhry and Gilleard, 2015). If the pre-existing mutations were the only source for the emergence of resistance, similar haplotypes would be present in the isolates (Chaudhry et al., 2016b), for example in the present study, 2 buffalo-derived isolates (CY321-B and CY322-B) carried four similar resistance haplotypes (BHA2-253S, BHA9-129G, BHA11-253S, BHA13-129G/253S). This striking similarity between the isolates provides evidence for the emergence of resistance by pre-existing mutations. If the recurrent mutations were the only source for the emergence of resistance mutations, different haplotypes would be present in the isolates (Chaudhry et al., 2016b), for example in the present study, 11 cattle-derived isolates have dramatically carried different haplotypes. This striking difference between the isolates provides evidence for the emergence of resistance by recurrent mutations.

In the present study, buffalo-derived 9 unique haplotypes of the 129G (GGC), 253S (TCT) resistance mutant and 129G (GGC)/253S (TCT) double resistance mutant were present in a single lineage, suggesting that there is a single origin of this mutation in *T. annulata* isolates examined. Similarly, 21 unique cattle-derived haplotypes of the 129G (GGC), 253S (TCT) and 262S (TCA) resistance mutant demonstrated evidence of a single origin. These results contrast with those of a previous study of the evolutionary origin of human protozoan parasite *P.vivax* dhfr resistance-conferring mutations demonstrated evidence of a single origin of pyrimethamine resistance (Shaukat et al., 2021).

In the present study, 129G (GGC), 253S (TCT) and 129G (GGC)/253S (TCT) and 262S (TCA) mutations were present in buffalo- and cattle-derived *T. annulata* collected from different livestock farms in the endemic regions and their phylogenetic relationship suggests that there is a high level of gene flow, possibly occuring due to unregulated animal movement. Notably, the two resistance haplotypes of the 129G (GGC) mutation predominate in 8 and 10 buffalo-derived *T. annulata* isolates and the other four resistance haplotypes of 129G (GGC), 253S (TCT) and 129G (GGC)/253S (TCT) mutations were present in 2 and 3 buffalo-derived *T. annulata* isolates respectively. In cattle-derived *T. annulata*, six resistance haplotypes of the 129G (GGC) mutation predominate between 7 to 10 different isolates and the other two resistance haplotypes of 129G (GGC) and 253S (TCT) mutations were present in 2 and 3 different isolates respectively. Overall, the unregulated animal movement has likely contributed to the spread of buparvaquone resistance haplotypes in the endemic region of Pakistan. The spread of resistance mutations may require a degree of reproductive isolation of the hosts and parasites with resistant alleles.

## 5. Conclusion

There are limited studies which have undertaken a genetic characteristic to detect buparvaquone resistance mutations in *T. annulata* of ruminant livestock under natural field conditions. Therefore, the data obtained in the present study provide a comprehensive analysis of whether cytochrome b loci are involved in the development of buparvaquone resistance and the first link between genetic mutations and drug resistance in *T. annulata*. There has been no study investigating the emergence of buparvaquone resistance in *T. annulata* and it is still unclear how resistance mutations spread in parasite populations under natural field conditions. The data then obtained in the present study provide information on single and or multiple emergences of resistance mutations and their spread models could have implications for novel buparvaquone resistance mitigation strategies.

## Acknowledgements

The study was financially supported by the Carnegie Trust Scotland and Biotechnology and Biological Sciences Research Council (BBSRC). Work at the Qaid-i-Azam University Pakistan and the University of Agriculture, Dera Ismail Khan Pakistan uses facilities funded by the Higher Education Commission of Pakistan.

---

**Supplementary Table S1.**
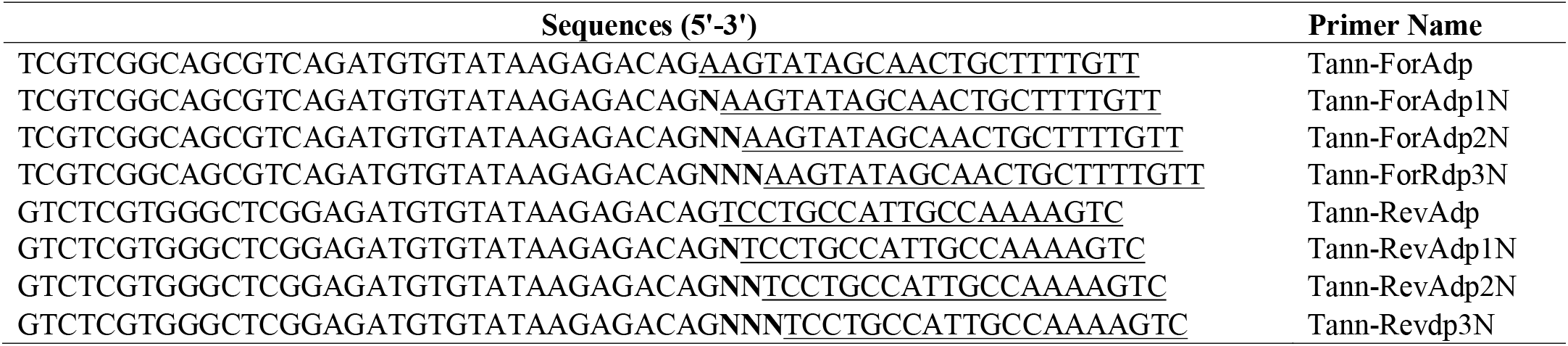
Primer sequences for Illumina MiSeq Library preparation. Tann-ForAdp/Tann-RevAdp primer sequence are underlined, N’s are in bold.

**Supplementary Table S2.**
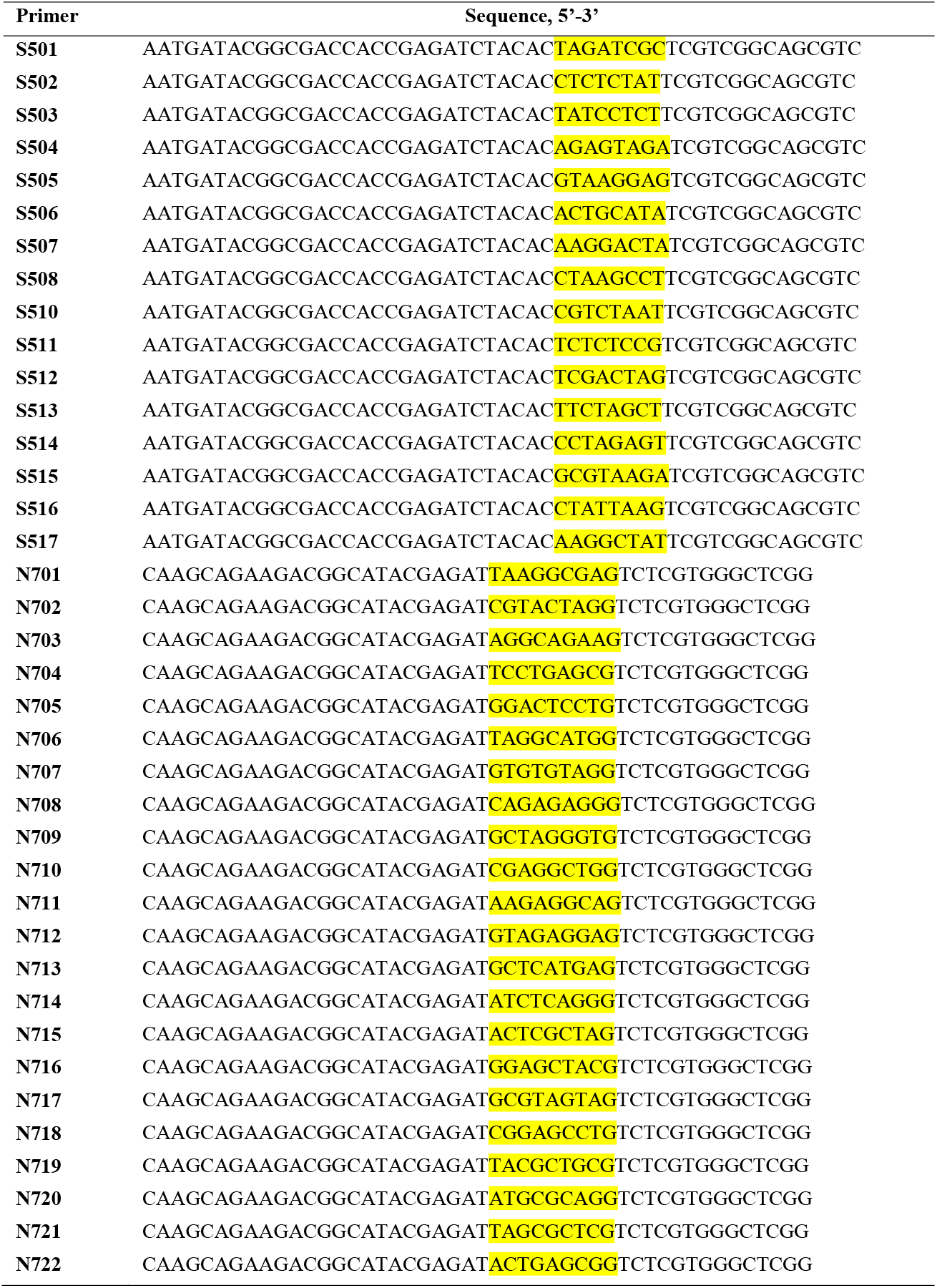
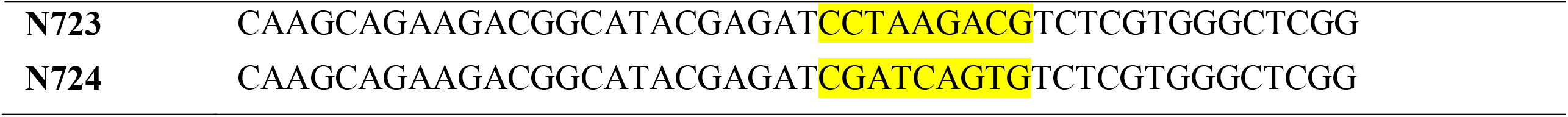
Sequences for forward and reverse barcoded primers (Nextera XT Index t v2). Index sequences are highlighted. Sequences from: Oligonucleotide sequences © 2018 umina, Inc. All rights reserved.

**Supplementary Table S3.**
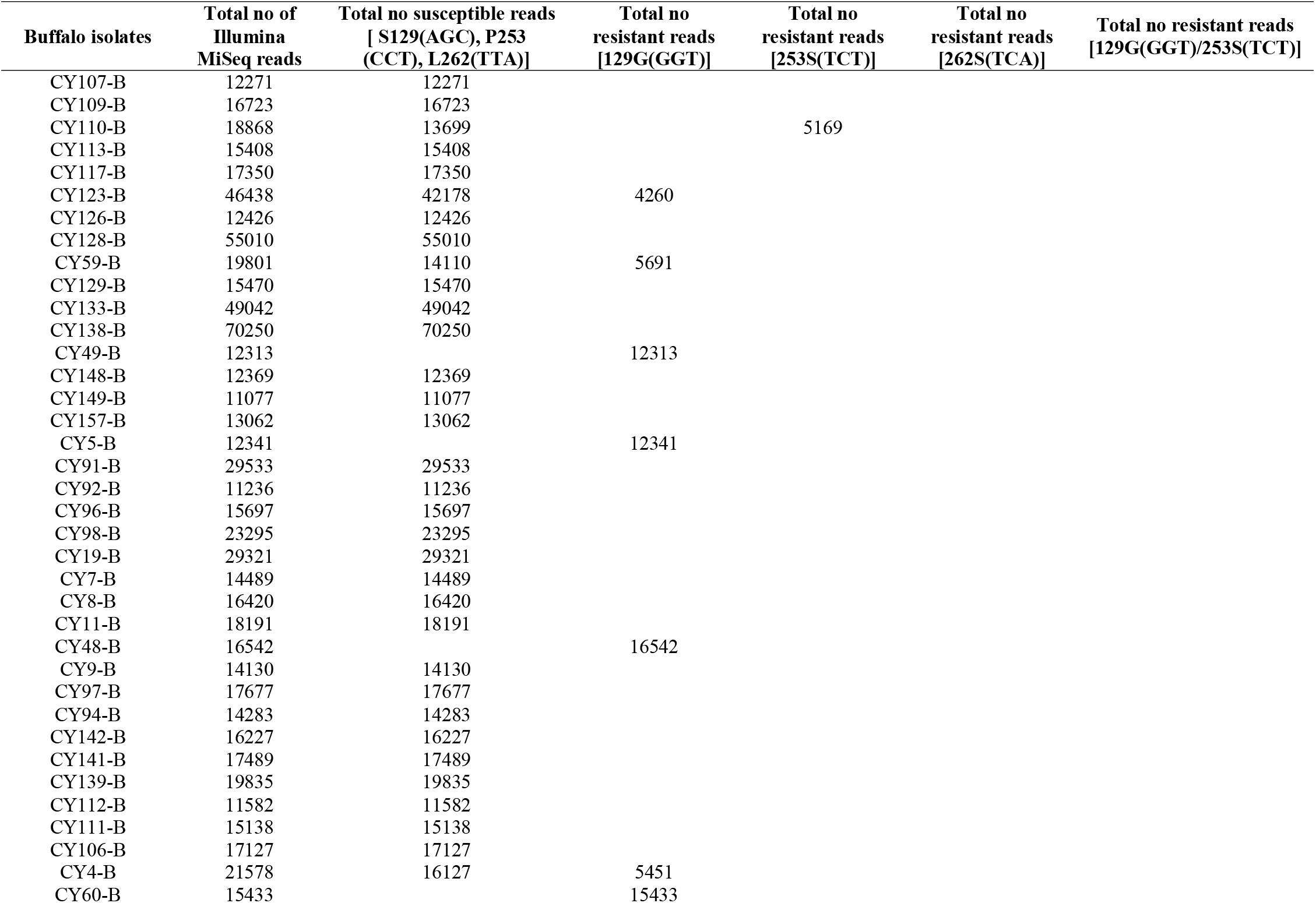
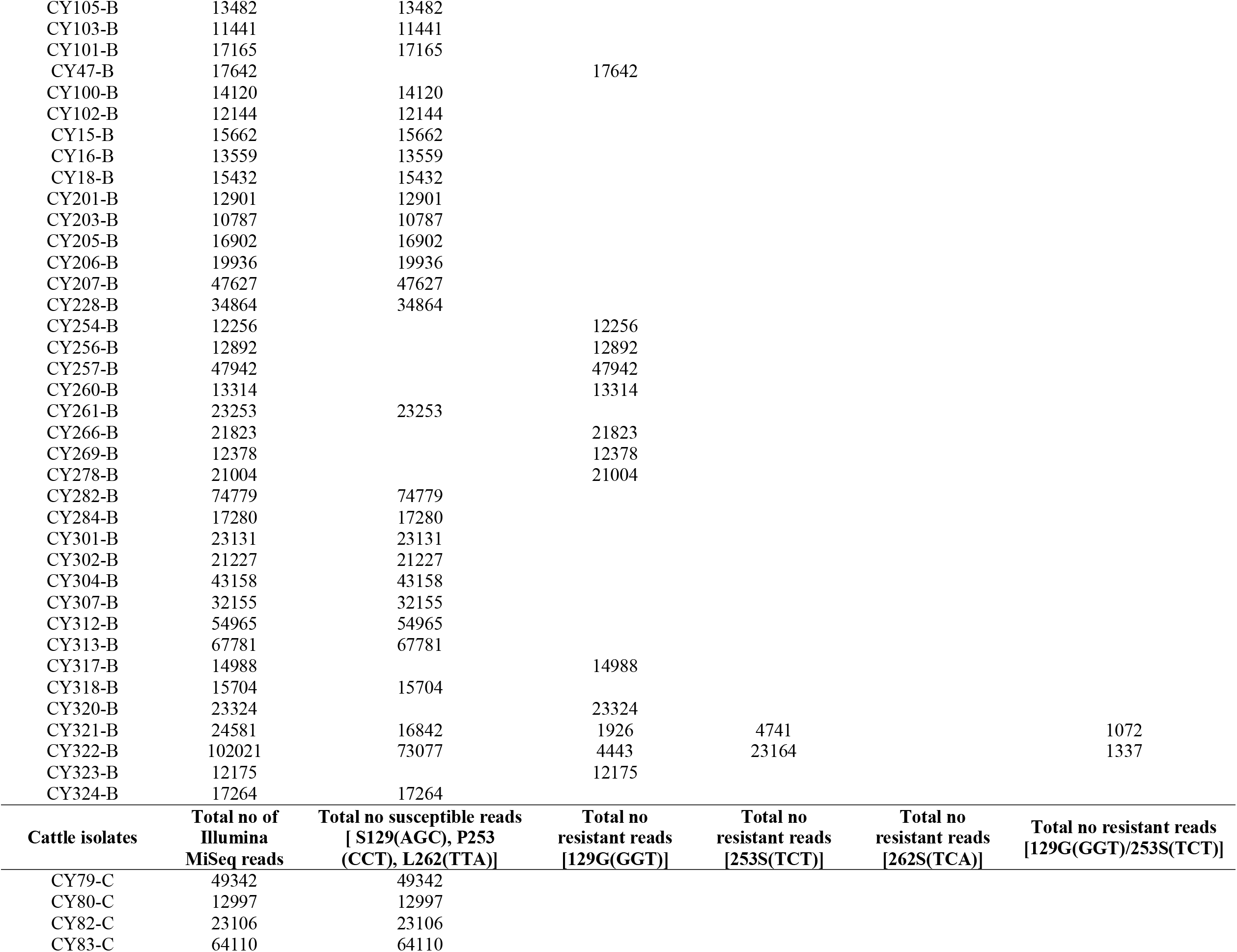
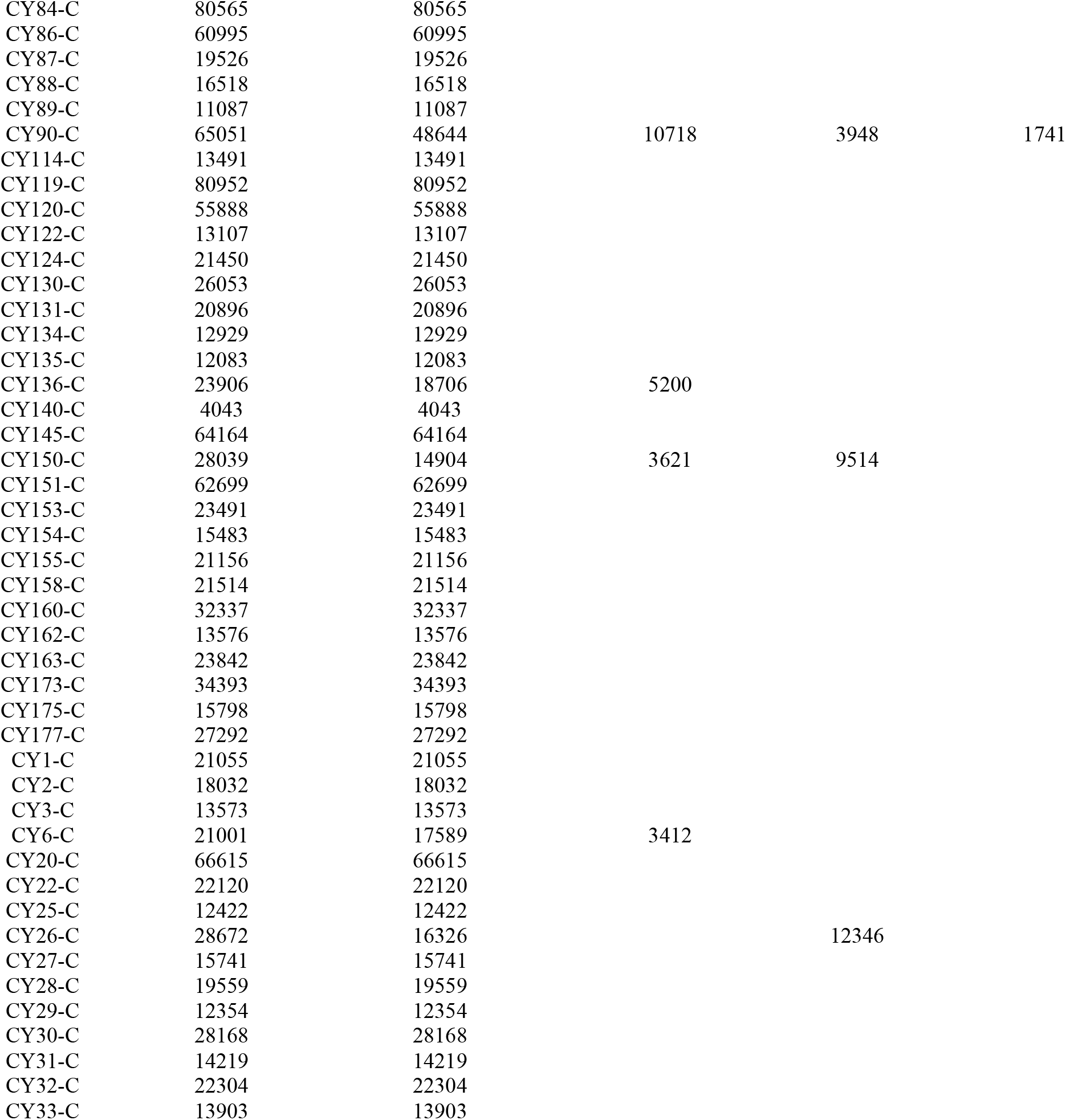
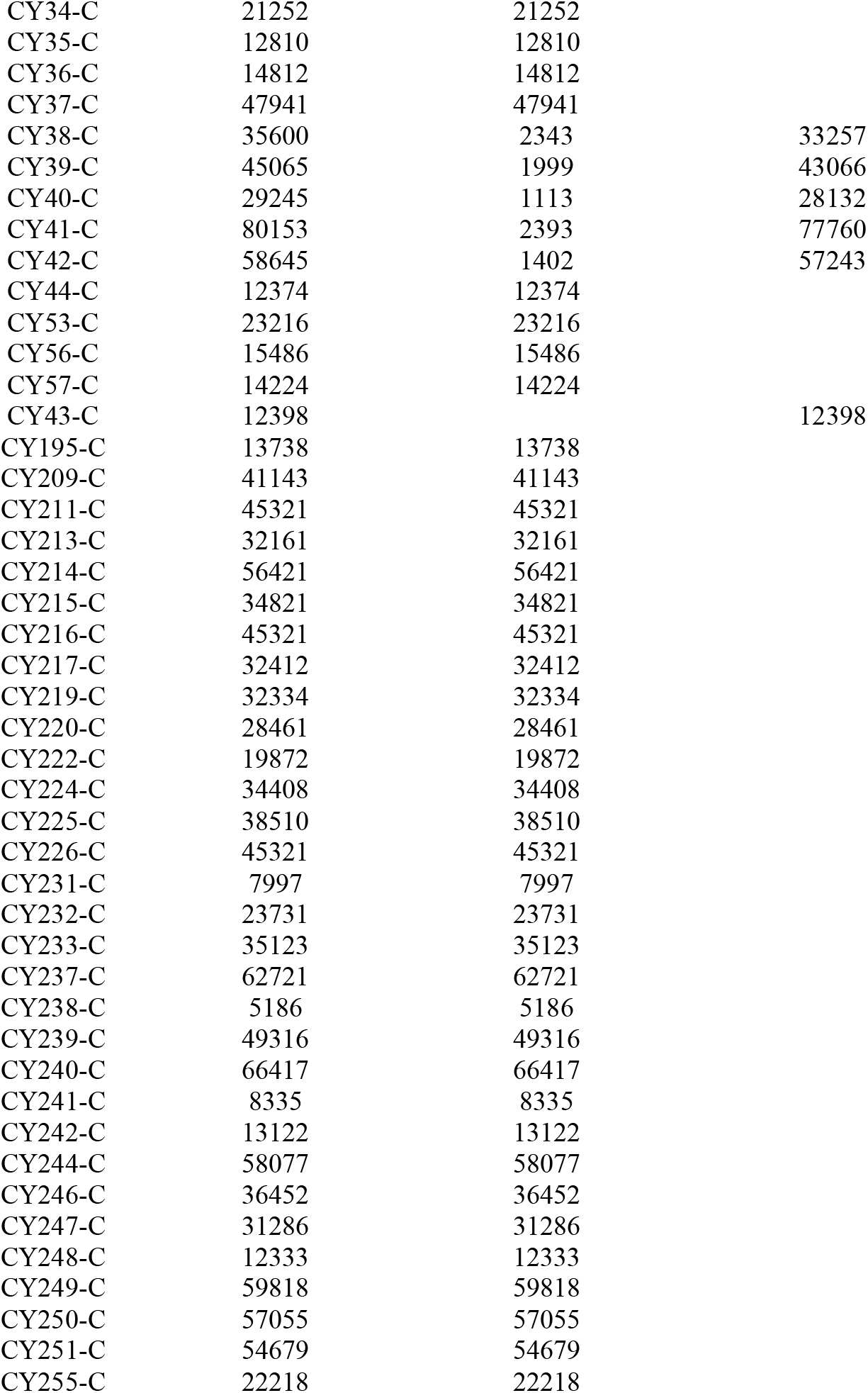
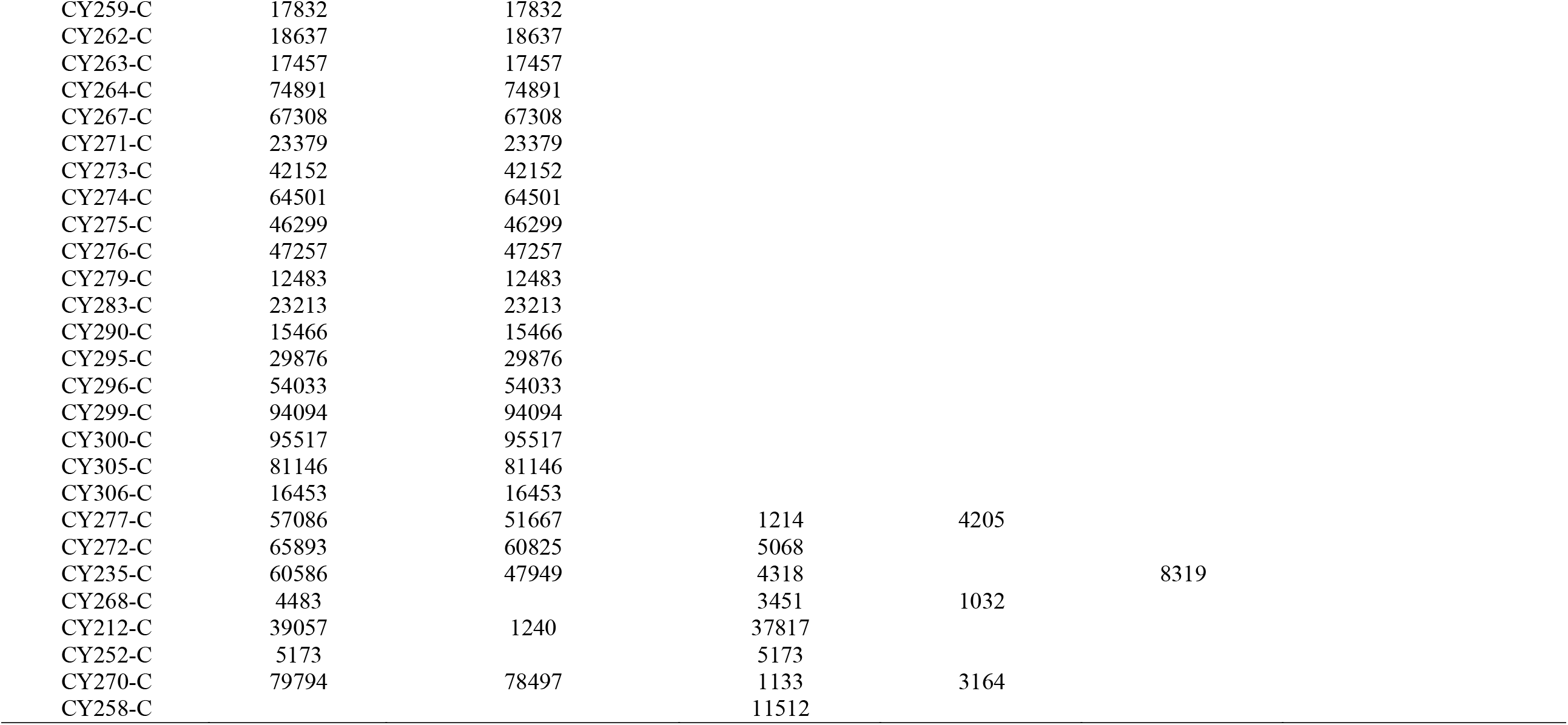
Deep amplicon sequencing data of the cytochrome b locus of *T. annulata* field samples. A total of 75 positive blood samples of buffalo and 119 from cattle were collected from veterinary clinics throughout the endemic regions of Pakistan.

## Notes

### Competing Interest Statement

The authors have declared no competing interest.

